# A Locally Produced Bioinoculant (H-Start) Increases Field Pea (*Pisum sativum*) Yield Using Reduced Fertilizer: A Pilot Study with Implications for Low-Input Agriculture

**DOI:** 10.64898/2026.10.19.760320

**Authors:** Ethan Done, Aminah Arif, Koenraad Duhem, Christopher Eskiw

## Abstract

Conventional agriculture inherently disrupts the natural soil microbiome. Although no-till practices reduce physical disturbance of the topsoil, nitrogen fertilizers and pesticides, particularly fungicides, still harm the soil food web. This weakens the soil’s capacity to support optimal biomass production and leaves large agricultural areas underproductive. Compost-based biostimulants (CBB) are heterogeneous mixtures of organisms from natural sources, such as compost, that can be added to soil to partially restore its natural microbiome. H-Start (HumaTerra Regen Ag Inc.) is one such CBB. Due to the lack of literature on H-Start, we aim to present a pilot study to identify potential effects of interest (α = 0.10) to inform future experimental designs. To achieve this, field pea (Pisum sativum cv. AAC Carver) was grown in Saskatchewan field soil in a growth chamber, with or without H-Start, at 75% or 100% of the recommended monoammonium phosphate (MAP) rate (n = 3 pots per treatment). By monitoring leaf size, leaf count, height, yield, and soil characteristics, we found that plants treated with H-Start were taller between days 21 and 28 (p = 0.089) and had a greater leaf size index (number of leaves x primary vein length^2^) over time (p = 0.002). H-Start at 75% MAP also produced the highest yield (p = 0.077). These results suggest H-Start influences the soil microbiome and the plants that depend on it. This is important because CBBs can be produced locally, which may make them increasingly relevant where conventional inputs are costly or difficult to obtain.

## 1. Introduction

Plants have the ability to convert carbon dioxide (CO_2_), water (H_2_O) and sunlight into usable metabolic energy (1). They often store this energy in carbohydrates (e.g., glucose, sugars and starch) that can act as an energy source for non-photosynthetic organisms (1). Humans have long propagated the most productive plants, and over time this process created various highly productive agricultural species that sustain ever-increasing populations. However, despite being able to generate energy from sunlight, plants are still dependent on soil for a range of chemicals, among which are nitrogen (N), phosphorus (P), potassium (K), calcium (Ca), magnesium (Mg), and sulphur (S) (2). It is important to note that the abiotic processes that cycle these nutrients are quite slow. Moreover, conventional agricultural practices remove large quantities of nutrient-laden biomass from soil while degrading the biotic pathways that enable replenishment. This has been formally recognized by the United Nations Food and Agriculture Organization in its World Soil Charter, which presents a series of nine principles that summarize the multi-faceted role the soil plays, and the threats to its ability to continue to serve these roles (3).

A way to act on these principles is to strengthen the symbiotic relationships plants have with the soil microbiome to support their growth (4,5,6). For example, *Pisum sativum* is capable of a process called nodulation, where the plant allows itself to become infected by select species of bacteria (for example, Rhizobium leguminosarum) (4,5). In exchange for carbon and a protected environment within the root nodule, these nitrogen-fixing bacteria use the enzyme nitrogenase to convert atmospheric nitrogen (N_2_), which plants cannot use directly, into ammonia that the plant can assimilate (4). Another example of this is arbuscular mycorrhizal fungi (6). These fungi increase a plant’s ability to take up various nutrients, including phosphorus and nitrogen, by colonizing the plant’s roots and trading soil-bound nutrients for sugars derived from photosynthesis (6). Arbuscular mycorrhizal fungi are also well documented to upregulate host defence genes in response to soil-borne pathogens (6). Currently, a limited diversity of these biological agents is often sold in the form of simple bioinoculants to increase crop yields. For example, as of 2026, one common bioinoculant is BASF’s Nodulator XL Peat, which claims to boost pea productivity by up to 8% using Rhizobium leguminosarum biovar viciae (strain 1435) (7). As promising as this is, evidence suggests that at least some of these bioinoculants require highly specific conditions to be effective (5). For example, a study by Kneen and LaRue (1984) found that *Pisum sativum* cultivar “Afghanistan” (derived from the Middle Eastern region) was unable to be colonized by *Rhizobium leguminosarum* strains isolated from distant regions (5). Because most biological processes function in a gradient rather than a binary fashion, it is worth questioning whether commercially available bioinoculants such as the aforementioned Nodulator XL Peat are effective in every geography in which they might be applied.

To address this, HumaTerra has developed a geographically localized bioinoculant called H-Start. H-Start is described by the company as “a targeted blend of a full spectrum of fungi and bacteria” (8). H-Start is produced by making compost that contains naturally occurring soil microbes that colonize the compost during the maturation phase (8, 9). Because H-Start contains a diverse community of strains from natural compost rather than a few carefully selected ones, the terminology Compost-Based Biostimulant (CBB) has been proposed (10). This contrasts with other bioinoculants such as Nodulator XL Peat that contain a single strain (7). HumaTerra uses DNA testing and advanced biochemical methods in the quality control of their products (11, 12, 13). The mixture is then compressed into a pellet that can be applied to a field. The fungi and bacteria survive the process, and H-Start thereby delivers intact viable microbes into the recipient soil (11). H-Start’s primary mode of action is hypothesized to proceed from complex interactions that occur during the coalescence of the recipient soil microbiome with a donor microbiome (H-Start), which results in a sequential carry-over of taxa, interplay among taxa, and a priming of functions in the recipient soil microbiome, which could account for mitigation or compensation of functions lost through various agricultural practices (14).

As such, we performed a small-scale pilot-level study to determine if H-Start is capable of (1) increasing yields and (2) reducing dependency on external fertilizer in *Pisum sativum* (cultivar AAC Carver) grown in Saskatchewan field soil. To achieve this, we applied H-Start at both 100% and 75% of the recommended Monoammonium Phosphate (MAP) fertilizer concentration. We then measured the height, number of leaves, primary-vein length, and yield. To confirm that our experimental setup allowed for interactions with the native soil microbiome, we used ITS amplicon sequencing on washed roots to ensure the soil contained active microorganisms, including potential pathogens such as *Fusarium* (Table 2). Through this research, we hope to promote and enable further studies into the potential benefits of locally produced compost-based biostimulants on agricultural crops.

## 2. Methods

### Planting

Twelve pots, labelled A-L, were filled with 1.000 kg ± 0.5% of soil sourced from Birch Hills, SK (Orthic Black Chernozem). This soil did not have a history of H-Start treatment. Pots were then divided into two equal groups for application of MAP fertilizer at 75% and 100% of the recommended rate of 112 mg/kg. All pots were hydrated with 250 mL of distilled water containing the appropriate dose of MAP. Three uniform and equally spaced insertions were made into each pot at a depth of 1 ¼ inches in a triangular arrangement and filled with one pea seed each.

Finely milled H-Start was made into a 560 mg/20 mL distilled water suspension, and 1.00 mL of this mixture was applied (pipetted) directly onto each embedded seed for the appropriate pots as shown in Table 1. Following this, pots were allowed to stand for one hour. Insertions were then covered with surrounding soil, and the pots were placed into a Conviron growth chamber with a 12-hour daytime temperature of 28.0 °C, a 12-hour nighttime temperature of 20.0 °C, and approximately 700 µmol/m^2^/s of light. Pots were placed in a random position within the growth chamber. Plants received a median of 100 mL of water per day. This heavy watering protocol was chosen to promote root rot in the plants, which is a common issue in Saskatchewan.

**Table 1:** Pot Treatment Array.

| Pots | H-Start | Fertilizer | Sprouted Plants |
|---|---|---|---|
| A-C | Yes | 75% | 7 |
| D-F | No | 75% | 9 |
| G-I | Yes | 100% | 9 |
| J-L | No | 100% | 9 |

Additional information regarding the pots is available in the supplementary data. Pot C contained 1 out of 3 plants.

### Root-Associated Microbial Community

A key part of how HumaTerra claims to work is by interacting with the native soil microbiome. As such, we wanted to ensure that the soil we used contained viable microorganisms and common pathogens. We thus harvested a plant from pots A, B, D, and E on day 49 and performed Nanopore ITS DNA sequencing on their washed roots individually (PathoScan Technologies Inc.), as shown in Table 2.

**Table 2:**
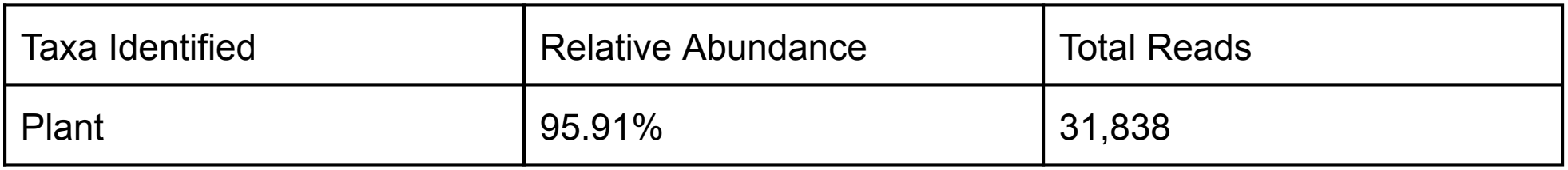
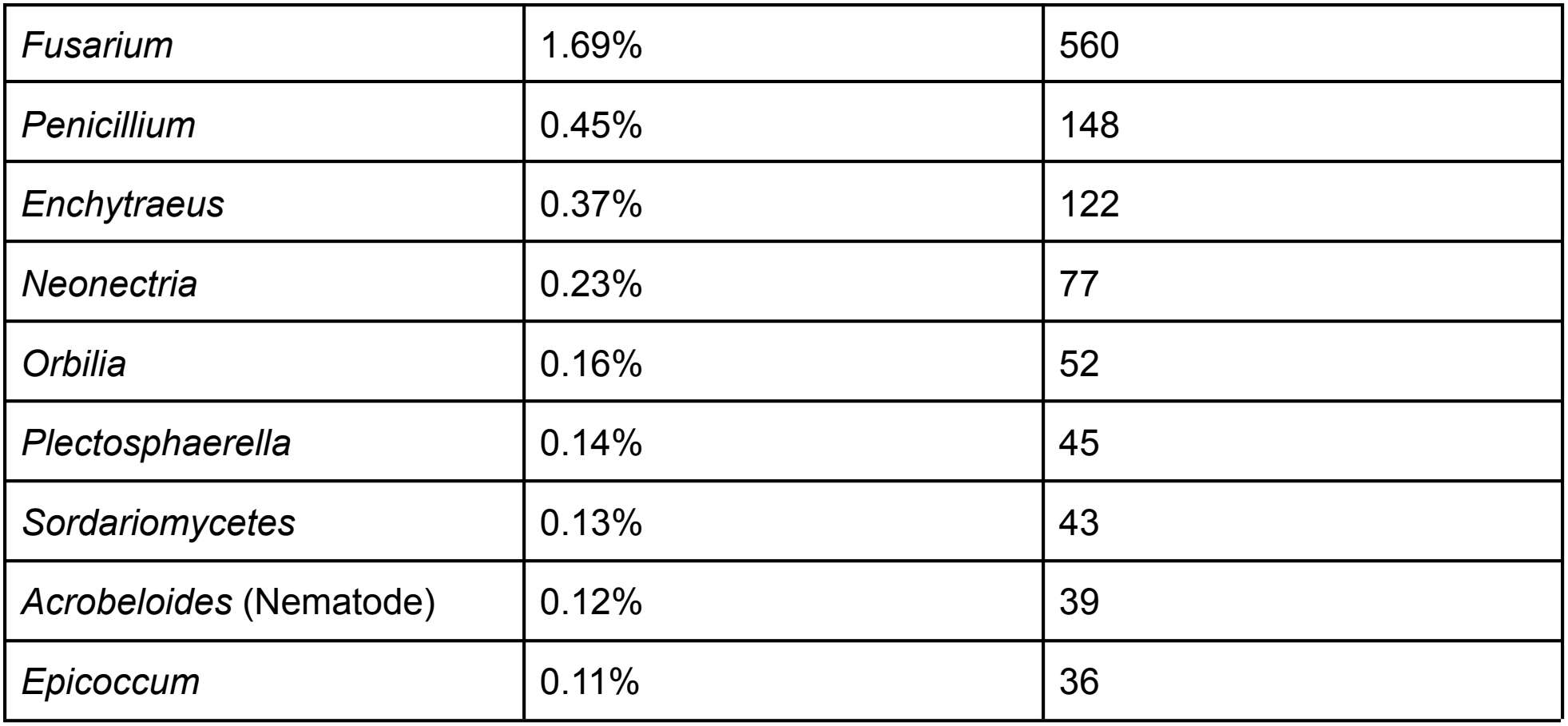
Top 10 Most Abundant Taxa Across All Samples.

### Measurement

To measure the plant height, a measuring stick was inserted into the soil on the first day of planting. Plant height thus represents the height of the plant above the soil line on the day observed. Leaves were counted manually. The primary-vein length was measured by measuring the length of the primary vein using a ruler on main-stem leaves counted down from the top of the plant. On each measurement day, the same leaf positions were measured on every plant; the positions sampled changed between days as plants grew. Peas were harvested after 132 days of growth. After harvest, peas were immediately removed from the plants and weighed. After 132 days of growth, the peas had dried on the plant.

We recognize that these methods are non-standard and that green coverage is a more commonly used statistic than primary-vein length. However, in this experiment, we wanted to elucidate variability at the plant level, and more common methods of measurement do not accurately reflect this. Following harvest, soil from each 100% MAP Pot (G - L) and a sample of the starting soil were homogenized and tested for pH, nitrogen, phosphorus, and potassium using a Lustre Leaf Rapid Soil Test Kit. Limited reagents prevented testing of the 75% MAP pots. N, P, and K were quantified at absorbance wavelengths of 540 nm, 710 nm, and 460 nm. Because testing reagents were limited, some samples were re-measured; where more than one reading was available, the latter reading was used. All original and repeated readings are reported in Supplementary Table 2. A calibration curve was used to determine how the colour of the pH test varied at different pH conditions, as shown in Figure 1. This curve, based on the ratio of A620 to A430, was then used to estimate the pH of the various soils. For ratios less than 0.877 (pH < 7), the equation pH = 4.122(A620/A430) + 3.387 was used. For ratios of 0.877 or above (pH ≥ 7), the equation pH = 2.071(A620/A430) + 5.185 was used.

**Figure 1:**
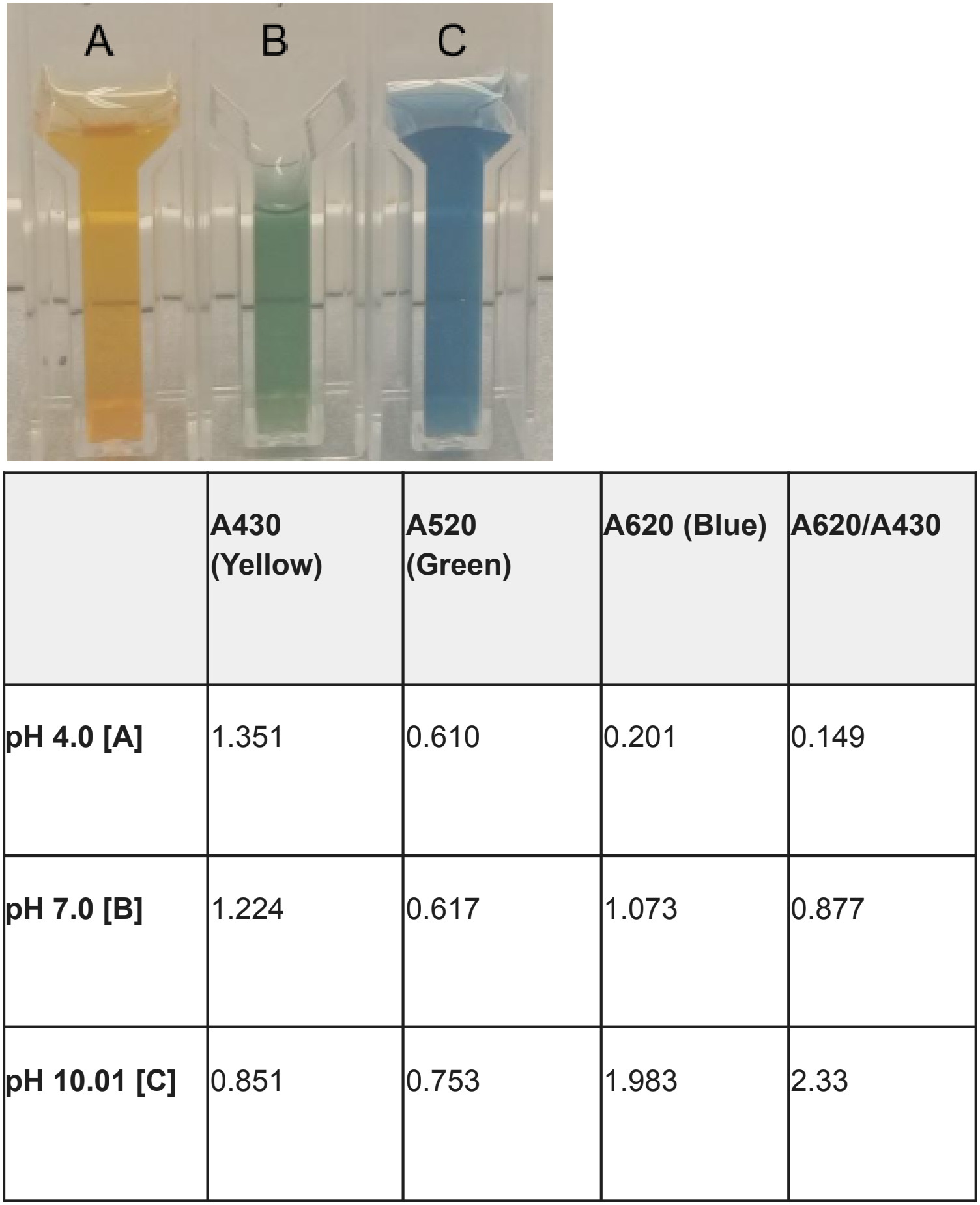
Calibration of pH Detection. Buffers of pH 4 [A], 7 [B], and 10.01 [C] were used to determine the colour of the pH testing kit.

### Statistical Analysis for Plant Height and Leaf Size Index

In this study, H-Start was applied to an equal number of pots with 75% fertilizer usage and 100% fertilizer usage. Because the primary aim of this study was to determine if H-Start is effective, and these are both concentrations that producers could use, samples were collapsed into an H-Start group and a no-H-Start group (n = 6) for analysis of plant growth. To evaluate the effect of the treatment groups on early plant growth (which is highly related to plant height), a Linear Mixed Model (LMM) was conducted (SPSS v29.1). The LMM approach was selected over a traditional repeated-measures ANOVA because it is robust to violations of sphericity (not measured due to small sample size) and allows explicit modelling of temporal correlations between repeated measurements on the same subjects. It can thus be considered akin to a regression for repeated measures data. Plant height (in mm) was entered as the continuous dependent variable. The collapsed treatment group (H-Start vs. No H-Start) and Time (days 14, 16, 19, 21, 24, 26, and 28) were entered into the model as fixed factors, alongside their interaction term (Treatment x Time). Plant heights were averaged within each pot, and because measurements were repeated on the same pots over time, pot was designated as the subject variable and Time as the repeated measure. A First-Order Autoregressive [AR(1)] covariance structure was specified for the residuals. This structure was chosen *a priori* as it appropriately assumes that successive measurements of plant height are more highly correlated with one another than measurements separated by longer intervals of time. Significant main effects and interactions were further investigated through the examination of Estimated Marginal Means (EMMs). When comparing main effects or specific time points between the two treatment groups, a Bonferroni adjustment was applied to all confidence intervals and p-values (however, with one comparison per time point, this did not alter the p-values). The “paste” option was used to inject the command “/EMMEANS = TABLES(Collapsed_Treatment*Days) COMPARE(Collapsed_Treatment) ADJ(BONFERRONI) in place of “/EMMEANS = TABLES(Collapsed_Treatment*Days)” to calculate *p*-values. Because this was a pilot study, all statistical tests were evaluated using an alpha level of 0.10.

Late-stage growth was quantified using a non-destructive leaf size index. Because leaves are approximately circular, leaf area scales with the square of primary-vein length; the index was therefore calculated for each plant as Index = L × V^2^, where L represents the total number of leaves and V represents the mean primary-vein length. These indices were aggregated at the pot level. A second LMM, utilizing the same fixed effects and AR(1) covariance structure as the height model, was employed to determine the effect of H-Start on leaf size index across the later growth stages (days 26, 35, 42, 49, 56, 63, 70, and 77). Statistical tests were evaluated using an alpha level of 0.10.

### Statistical Analysis for Yield Data

Yield data were not independent due to the fact that multiple peas were grown per pot, and as such, effects were observed at the pot level. To account for attrition throughout the experiment, the weight of peas collected per pot was divided by the number of plants at harvest in that pot. Pea weight data were continuous. Due to the low sample size (n = 3 per group), normality could not be assessed using pots as the unit of replication. However, all treatment groups were normally distributed at the individual plant level (1-Sample Kolmogorov-Smirnov Test, 0.196 < Z < 0.254, *p* = 0.200) and, combined with the tolerance of ANOVAs to non-normal distributions, this was deemed suitable for this pilot study. All treatments also had equal variance (Levene’s Test, F_3, 8_ = 2.053, *p* = 0.185). As such, to determine if the combination of H-Start and Fertilizer had an effect on the yield of pea plants, a 1-Way ANOVA was performed at alpha = 0.10 (SPSS v29.1). A Fisher’s Least Significant Difference (LSD) post hoc test was used for visualization to indicate the potential for further study.

To determine the required number of replicates for future studies, a power analysis was performed. Given the claims made by BASF about their Nodulator XL Peat, an 8% effect size (Δ) was assumed. This was determined to be 0.052 g for the No H-Start, 75% MAP group. This value was used conservatively because it represents the smaller of the two No H-Start hypothetical effect sizes. Standard deviation (σ) was calculated to be 0.98 (H-Start, 75% MAP, n = 5), 0.60 (No H-Start, 75% MAP, n = 6), 0.50 (H-Start, 100% MAP, n = 9), and 0.61 (No H-Start, 100% MAP, n = 9). These were weighted by sample size to create a representative standard deviation of 0.64. The critical significance value used was 1.96 (Z_a/2_). The critical value for power used was 0.84 (Z_b_). Together, these values represent an experiment with a 95% probability of not having a false positive (alpha) and an 80% probability of finding an effect when there is one (power). This differs from the *α* = 0.10 threshold used in this study because a future study would not be at a pilot scale. The required sample size was calculated using the formula n = 2 x (Z_a/2_ + Z_b_)^2^ x σ^2^ / Δ^2^.

### Statistical Analysis for Nitrogen Testing

Due to the low replicate value for the nutrient testing (n = 3 for H-Start vs. No H-Start), and a lack of a calibration curve capable of mapping absorbance to nutrient level, we assumed that the test values were normally distributed despite there being too few replicates to run a 1-Sample Kolmogorov-Smirnov Test. We then performed a Levene’s test to ensure homoscedasticity between the H-Start and No H-Start pots (Levene’s Test, F_4_ = 0.090, *p* = 0.779). As such, an Independent Samples T-Test was performed to compare the Nitrogen nutrient test absorbance values of pots G, H, and I (H-Start + 100% MAP) with pots J, K, and L (100% MAP).

## 3. Results

### Early Plant Growth

To begin, we found that there was a significant interaction between H-Start and Time of plant growth on plant height (Linear Mixed Effects, F_6, 59.86_ = 1.939, *p* = 0.089). Early in plant growth (days 14, 16, and 19), there is no difference in plant height between treatment groups (p = 0.265, p = 0.167, and p = 0.108, respectively). However, by day 21, plants with H-Start were an average of 23 mm taller than plants without H-Start (*p* = 0.044). This trend held on days 24 and 26 (*p* = 0.026 and *p* = 0.018, respectively). By day 28, peas with H-Start were an average of 32 mm taller than plants without it (*p* = 0.011), as shown in Figure 2. It must be noted that this result matches what would be expected for a bioinoculant, as early pea growth is driven primarily by stored nutrients, which are independent of external factors.

**Figure 2:**
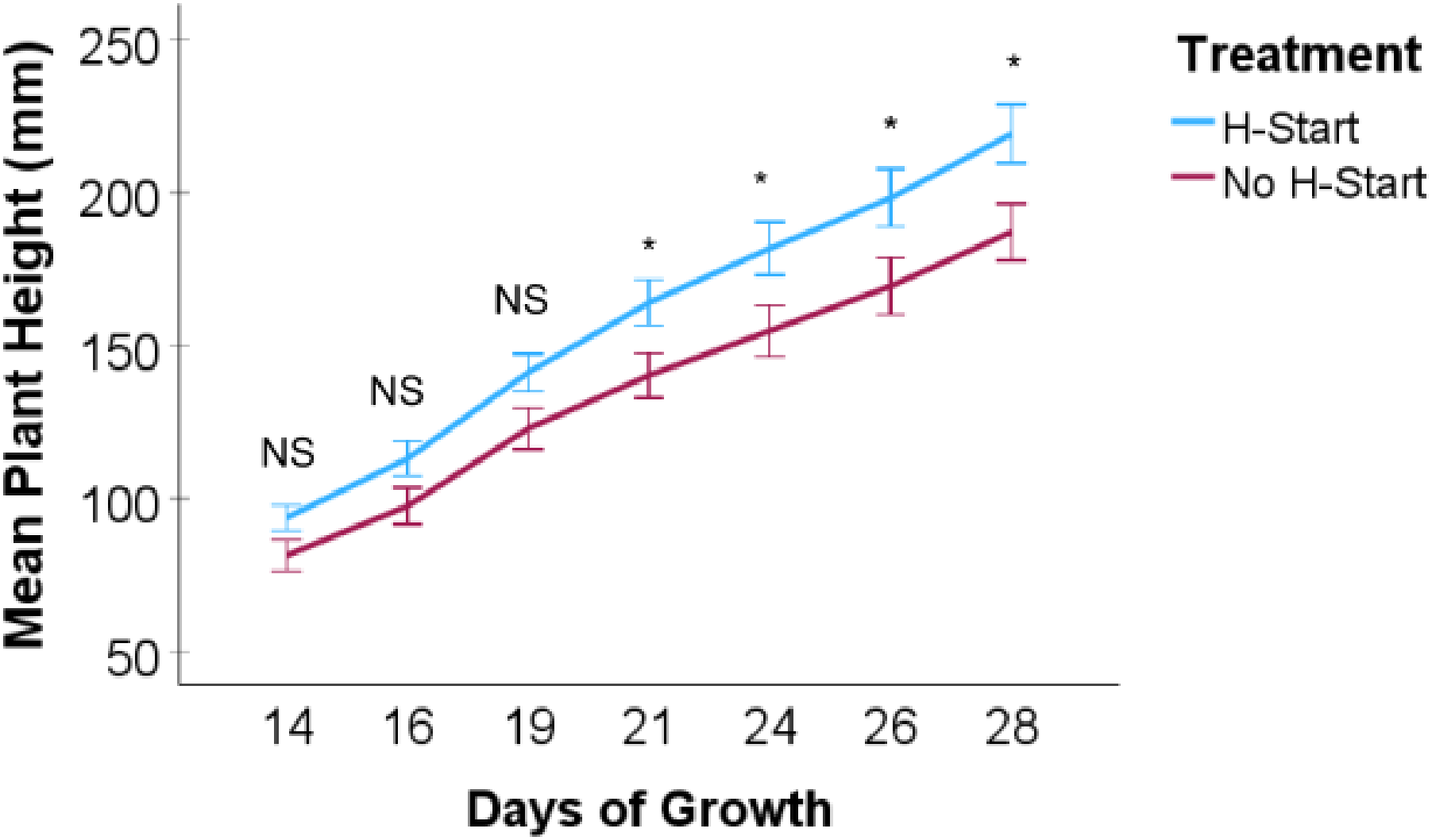
Effects of H-Start on Plant Growth. Pots of pea plants (n = 6 per group) were grown with or without the bioinoculant H-Start and with varying levels of MAP fertilizer. Data points represent the mean height of plants (mm). Error bars represent the standard error of the mean. NS represents a lack of significance at alpha = 0.10. * Represents significance at alpha = 0.10.

### Late Plant Growth

As plants grow, they begin to put increasing energy into creating photosynthetic surface area rather than height. As such, the leaf size index was used as a proxy for later plant growth. Interestingly, we again found that there was a significant interaction between H-Start and Time of plant growth on leaf size index (Linear Mixed Effects, F_7, 66.19_ = 3.712, *p* = 0.002). Indeed, we find that on days 26, 35, 42, and 49, both H-Start-treated plants and non-H-Start-treated plants have the same mean leaf size index (*p* = 0.675, *p* = 0.461, *p* = 0.205, *p* = 0.121). However, at days 56 and 63, the H-Start plants had leaf size indices 2455 mm^2^ and 3069 mm^2^ more than the non-H-Start plants (*p* = 0.094 and *p* = 0.034, respectively). This reverted to non-significance at day 70 and day 77 (*p* = 0.782, *p* = 0.121). It is worth noting that until day 77, H-Start plants consistently had a higher leaf size index than non-H-Start plants, as shown in Figure 3.

**Figure 3:**
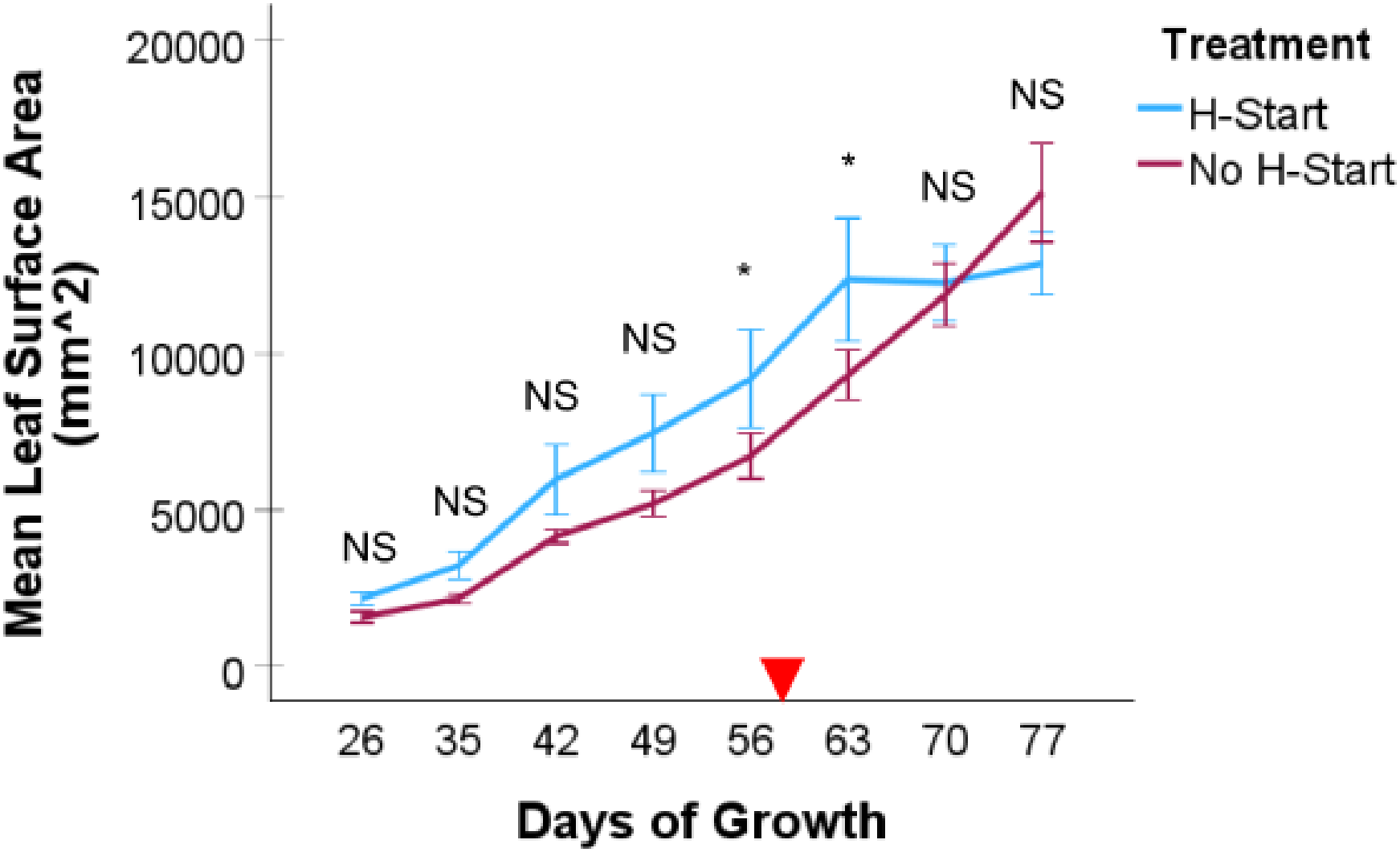
Effects of H-Start on Leaf Size Index. Pots of pea plants (n = 6 per group) were grown with or without the bioinoculant H-Start and with varying levels of MAP fertilizer. Data points represent the mean leaf size index of plants (mm^2^). Error bars represent the standard error of the mean. NS represents a lack of significance at alpha = 0.10. * Represents significance at alpha = 0.10. The red triangle represents the first observation of flowering.

### Yield Data

Using the pilot-study significance threshold of *α* = 0.10, we found that H-Start had a significant effect on yield when 75% MAP was applied (1-Way ANOVA, F_3, 8_ = 3.331, *p* = 0.077), as shown in Figure 4.

**Figure 4:**
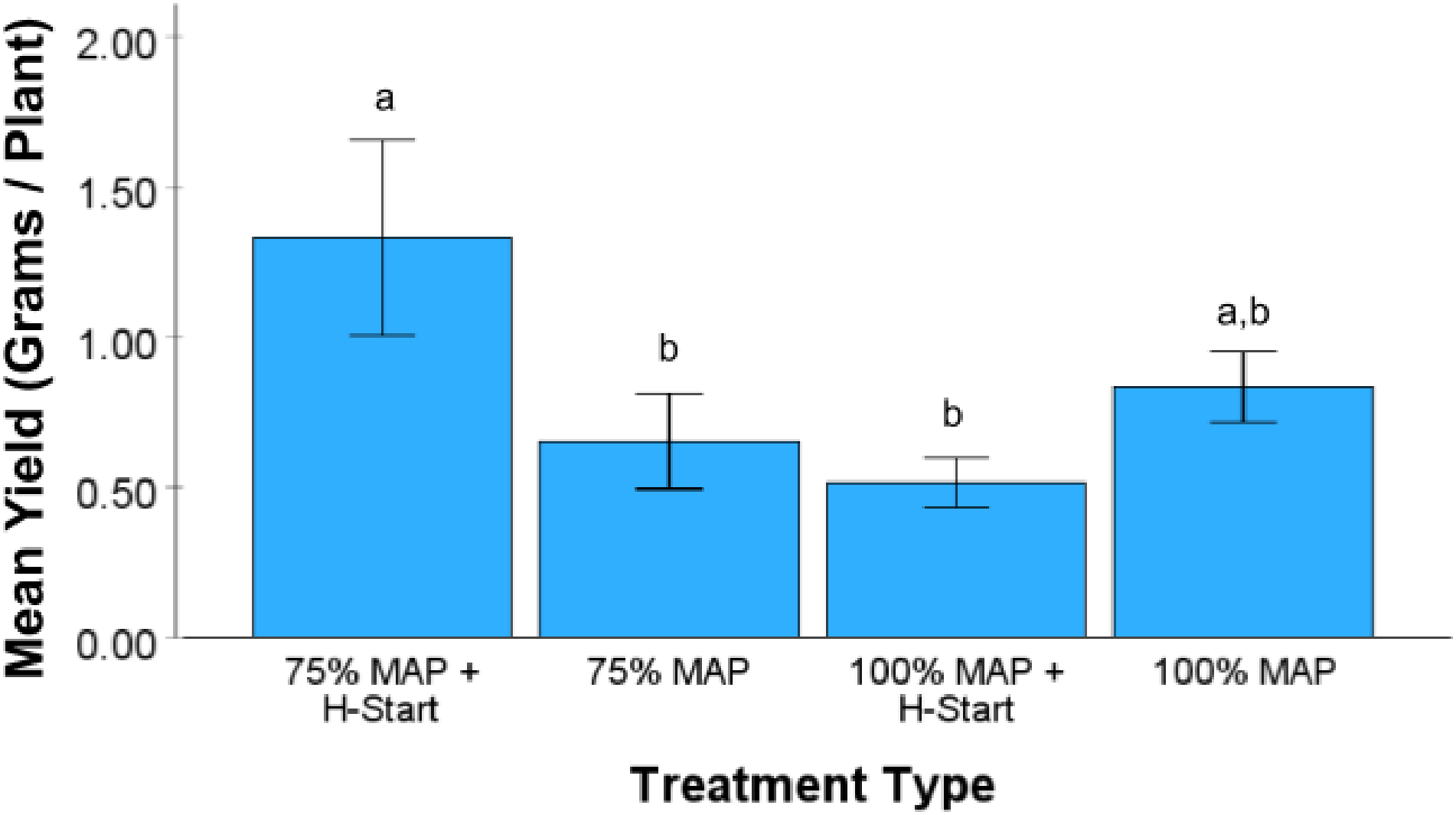
Mean Pot-Level Yield of Various Treatments. Pots of pea plants (n = 3 per group) were grown with various combinations of the bioinoculant H-Start and MAP fertilizer. Bars represent the mean pea yield per plant. To account for experimental attrition, individual plant yields were first averaged within each pot before calculating the overall treatment means. Error bars denote the standard error of the mean. Different letters denote statistical significance using Fisher’s LSD post-hoc. Pea weights are shown in Supplementary Table 1. Pot C contained a single plant (1.945 g); the highest-yielding individual plant in this group was in Pot B (2.444 g; Supplementary Table 1). Because of this, Pot C was not excluded.

It must be noted that the experiments contained herein are exploratory; hence, the *α* = 0.10 threshold was used. However, using a technique called a power analysis, we determined that future studies wanting to conclusively test if H-Start increases yield will likely want to include at least 2,376 independent plants per treatment group if significance is determined at alpha = 0.05. This represents approximately 150 ft^2^ of space, assuming a 3-inch planting distance, something that is easily achievable in a field setting.

### Post-Plant Soil Testing

To begin, soils were tested for pH to determine the effect of microbial activity from H-Start. As shown in Table 3, H-Start seems to increase the variability of the soil pH, with both the highest and lowest pH being found in this group (range = 3.02). This is in contrast to the MAP fertilizer-only group (range = 0.82). We note here that this may be due to the limited number of pots used in this trial and, as such, further research should be conducted to confirm this finding. We also note that Pot I’s absorbance reading exceeded the range of the pH standards and the reliable range of the spectrophotometer, so its pH estimate is uncertain. However, it is clearly unique among pots, leading to our hypothesis that H-Start causes increased variability.

**Table 3:** Pot-Level Soil pH Readings.

| **Pot** | **Condition** | **A620** | **A430** | **A620/A430** | **Final pH** |
|---|---|---|---|---|---|
| **Starting Soil** | NA | 0.606 | 1.305 | 0.4644 | **5.30** |
| **G** | H-Start + 100% MAP | 0.830 | 1.930 | 0.4301 | **5.16** |
| **H** | H-Start + 100% MAP | 0.728 | 1.246 | 0.5843 | **5.80** |
| **I** | H-Start + 100% MAP | 3.234 | 2.234 | 1.4476 | **8.18** |
| **J** | 100% MAP | 1.960 | 1.798 | 1.0901 | **7.44** |
| **K** | 100% MAP | 0.944 | 1.113 | 0.8482 | **6.88** |
| **L** | 100% MAP | 0.634 | 0.807 | 0.7856 | **6.63** |

Next, we analyzed the soil concentration of N, P, and K as shown in Figure 5. Nitrogen levels were of particular note, as there seemed to be a difference between pots with H-Start and pots without. We thus analyzed these and found that while the effect of H-Start was not statistically significant (Independent Samples t-test, t_4_ = 2.009, *p* = 0.115), the absorbance reading from the nitrogen test was 61% higher in these pots compared to pots without H-Start. Both re-measured nitrogen samples (pots H and L) shifted in favour of the H-Start group. Using only the original measurements, H-Start pots had a 31% increase in nitrogen absorbance compared to non-H-Start treated pots.

**Figure 5:**
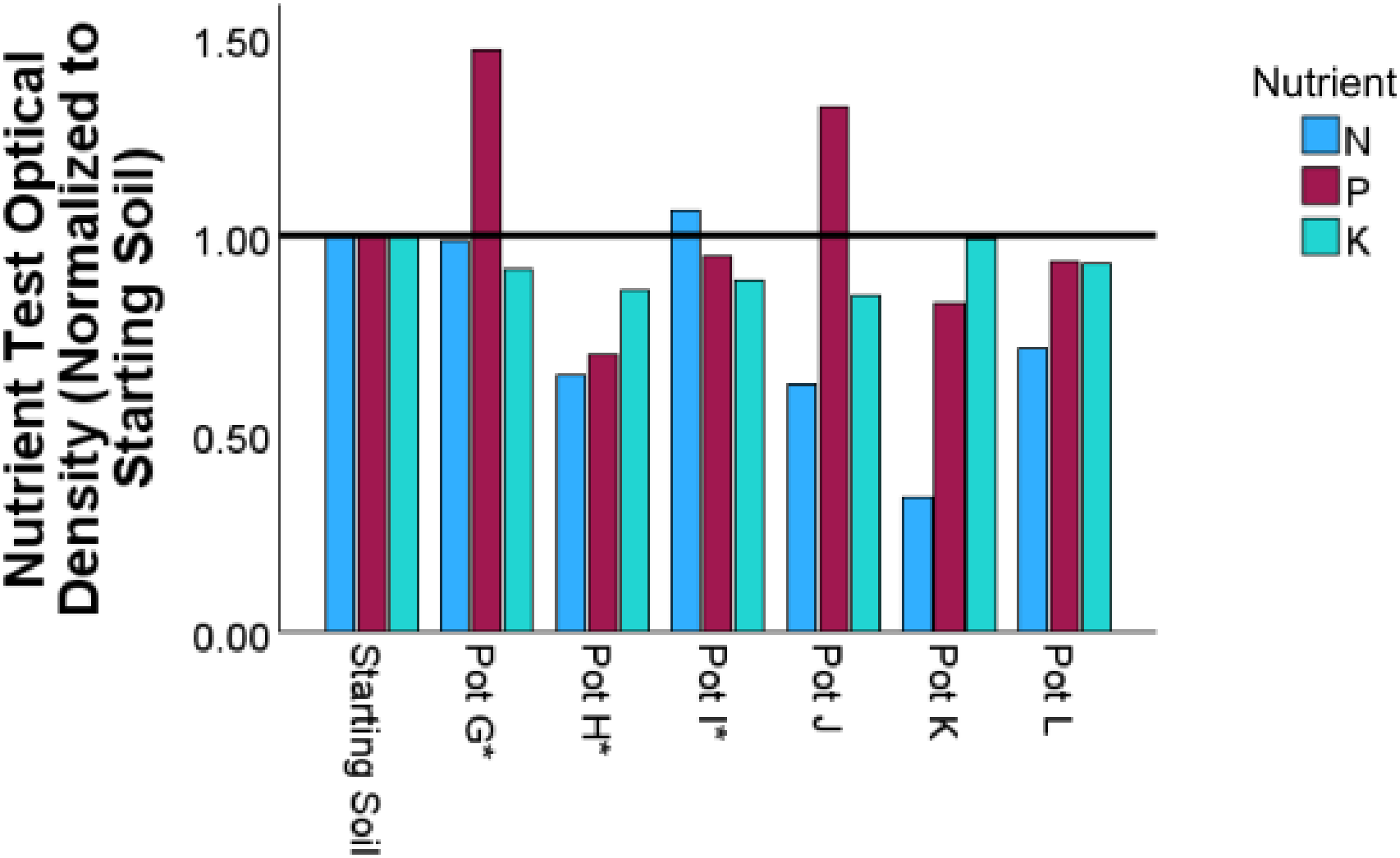
Nutrient Testing Results. A Lustre Leaf test kit was used to determine the N, P, and K levels of soil after pea plant growth. Starting soil is the soil used to grow the plants. All other pots had 100% of the recommended dose of MAP fertilizer added. H-Start was added to pots G, H, and I as denoted by an asterisk. Bars represent the optical density of the nutrient testing kit, with higher optical density corresponding to a higher nutrient composition. All values are normalized to the starting soil for comparison. The black bar denotes no change from the starting soil.

## 4. Discussion

This study represents an evaluation of the effects of the compost-based biostimulant H-Start in field peas (*Pisum sativum*). Using an exploratory significance threshold of α = 0.10, we found H-Start altered plant height over time (p = 0.089), with H-Start plants being 32 mm taller by day 28 (p = 0.011). H-Start also increased the leaf size index over time (p = 0.002). Finally, when paired with 75% of the recommended dosage of MAP fertilizer, H-Start increased yield (1-Way ANOVA, F3,8 = 3.331, p = 0.077; Fisher’s LSD). H-Start did not significantly change soil nitrogen, although nitrogen test readings were approximately 61% higher in pots with H-Start than in pots without it.

We note that the results contained herein are not definitive. Based on the variability we observed, we recommend that future experiments contain approximately 150 ft^2^ of plants per treatment group so that all of H-Start’s effects, including on soil chemistry, can be observed with increased accuracy. We also note that H-Start is a compost-based biostimulant whose effect is believed to be largely dependent on microbes, which are notoriously heterogeneous and can experience evolution during the manufacturing process. As such, we recognize that a limitation of this study is the use of only a single batch of H-Start. Finally, we would also like to note that this study suggests that H-Start is more effective in plants exposed to 75% of the recommended rate of MAP than 100% of the recommended rate. We note that this may be a true biological effect due, for example, to MAP possibly being toxic to the microbes within H-Start. However, we would like to express caution when interpreting this result, as the study contained herein is likely low-powered. As such, it may also be that, by chance, the seeds contained in the 100% MAP treatment group responded less efficiently to H-Start. We also believe that the soil chemistry tests suffered from the same low-power issue. As such, we encourage the results to be interpreted as *bona fide* signals only when they pass the standard threshold of *p* = 0.05, suggestive signals only when they pass *p* = 0.10, and as inconclusive at *p* > 0.10 rather than negative. We also did not include a sterilized H-Start control, so the effects of its microbes cannot be separated from those of its organic matter and nutrients. We also note that plant numbers differed between pots and that soil chemistry was measured only in the 100% MAP pots.

Nevertheless, it should be noted that H-Start’s use of microbes (or at least, microbe-derived factors created during the compost maturation process) as its primary mode of action makes it attractive as an input option for low- and middle-income countries (LMICs). This is because it allows HumaTerra’s proprietary H-Start manufacturing process to remain relatively straightforward by requiring no complex inputs, such as large amounts of electricity, that cannot be sourced locally. In locations that are unable to procure traditional agricultural inputs due to logistics, costs, or other factors, we believe that this localized approach to production could be greatly beneficial. As such, we believe H-Start, or other locally manufactured derivatives, offer a promising alternative to the traditional agri-chemical supply chain and encourage the use of the data contained herein to guide the experimental design of future investigations evaluating the effects of H-Start or other locally produced bioinoculants.

## 5. Author Statements and Disclaimers

### Author Contributions

E. Done designed and managed the experiment, collected and analyzed the data, wrote the manuscript, acquired funding, and arranged DNA sequencing. A. Arif assisted with experimental execution. K. Duhem provided input on the experimental design, contributed background information on H-Start, and reviewed the manuscript. C. Eskiw supervised the project.

### Data Availability

The data supporting this study are provided in the Supplementary Material or are available from the corresponding author upon reasonable request.

